# Acquired Amphotericin B Resistance Attributed to a Mutated *ERG3* in *Candidozyma auris*

**DOI:** 10.1101/2025.03.30.646105

**Authors:** Lauryn Massic, Laura A Doorley, Sarah J Jones, Irene Richardson, Danielle Denise Siao, Lauren Siao, Philip Dykema, Chi Hua, Emily Schneider, Christina A. Cuomo, P. David Rogers, Stephanie Van Hooser, Josie E Parker, Steven L Kelly, David Hess, Jeffrey M Rybak, Mark Pandori

## Abstract

First identified in 2009, *Candidozyma auris* (formerly *Candida auris*) is an emerging multidrug resistant fungus that can cause invasive infections with a crude mortality rate ranging from 30-60%. Currently, 30-50% of *C. auris* isolates are intrinsically resistant to amphotericin B. In this work, we characterized a clinical case of acquired amphotericin B resistance using whole genome sequencing, a large-scale phenotypic screen, comprehensive sterol profiling, and genotypic reversion using CRISPR. Data obtained in this work provides evidence that a deletion resulting in a frameshift in *ERG3* contributes to the observed resistant phenotype.

Characterization of this isolate also revealed a fitness cost is associated with the abrogation of ergosterol production and its replacement with other late-stage sterols. This article presents a clinical case description of amphotericin B resistance from a frameshift mutation in *ERG3* in *C. auris* and marks an advancement in the understanding of antifungal resistance in this fungal pathogen.

## Introduction

In 2019, the Centers for Disease Control and Prevention (CDC) designated *Candidozyma auris* (previously known as *Candida auris*) as an urgent antimicrobial threat making it the first fungal pathogen to be raised to this level of concern (1, 2). This is attributed to the multidrug resistant characteristics of *C. auris*, its capacity to spread easily in health care facilities, and its potential to cause invasive candidiasis especially among patients with weakened immune systems (3-6). Out of the three primary antifungal classes approved for the treatment of *Candida* infections, approximately 80-93% of *C. auris* isolates are resistant to fluconazole (triazole drug class), 35-50% display resistance to amphotericin B (polyene drug class), and 5-7% show resistance to the echinocandin antifungals (7-9).

Amphotericin B is frequently utilized due to its broad-spectrum of activity and success in treating systemic fungal infections (10). At the time of manuscript preparation, there are no clinical amphotericin B susceptibility breakpoints set forth by the Clinical Laboratory Standards Institute (CLSI) for *C. auris* (11), however the CDC has established tentative amphotericin B minimum inhibitory concentration (MIC) breakpoints for *C. auris* at 2 µg/mL. However, up to a third of *C. auris* isolates from the United States have MICs of 1 µg/mL thus caution is urged for *C. auris* amphotericin B MIC interpretation (11-14).

Even though amphotericin B has been used in clinical practice since the 1950s very few mechanisms of drug resistance have been documented. Previously, acquired cases of amphotericin B resistance in *Candida spp*. have been shown to be associated with mutations in the genes encoding the ergosterol biosynthesis pathway (8). In *C. auris*, the only established mechanism of clinically acquired amphotericin B resistance is attributed to an indel in *ERG6* (15). Additionally, a nonsense mutation in *ERG3* has once been associated with amphotericin B resistance in a single clinical isolate but has yet to be tested/confirmed (16). Other mechanisms of amphotericin B resistance have been identified *in vitro*. Amphotericin B resistance can be induced by culturing *C. auris* in the presence of sub-lethal doses of amphotericin B (17). In other *Candida spp*., mutations in *ERG2, ERG3, ERG4, ERG5, ERG6*, and *ERG11* have been found to be associated with reduced susceptibility to amphotericin B (8, 10, 18-20).

Here, we present a case of acquired clinical amphotericin B resistance which is attributed to a mutation of the *C. auris ERG3* gene causing a premature stop codon in the C-5 sterol desaturase. We further investigated this isolate through a phenotypic screen, CRISPR-Cas9 repair, and sterol profiling. Our results support a causal link between the *ERG3* mutation and much of the amphotericin B resistance phenotype.

## Methods

### Collection, Culturing, and Confirmation of Specimens

The first isolate (LNV001) from a patient was collected from the bronchial lavage in July 2022 while the second isolate (LNV002) from the same patient was collected from the urine in September 2022. Both samples were initially grown in Salt Sabouraund Dulcitol broth at 38 °C at 250 rpm for 24-48 hours. From culture, a 10 µL loop was used to streak culture on CHROMagar™ *Candida* (CHROMagar™, Paris, France) where it was grown for 24-48 hours at 36 °C. Plates displayed a pinkish purplish growth and were confirmed as *C. auris* by matrix-assisted laser desorption ionization–time of flight mass spectrometry (MALDI-ToF) with reference library MALDI Biotyper CA library (version 2022) (Bruker, Billerica, MA).

### DNA Extraction

Clinical isolates underwent bead-beating (FastPrep-24, MP Biomedicals, Irvine, CA) for 4 cycles at 6.0 m/sec for 30 seconds with 5-minute pauses in between. Then genomic DNA (gDNA) from isolates was isolated using PureFood Pathogen Kit on the Maxwell RSC (Promega, Madison, WI) per manufacturer’s protocol.

### Library Prep and Whole Genome Sequencing

Extracted gDNA was library prepped using DNA Prep Kit (Illumina, San Diego, CA) per manufacturer’s recommended protocol utilizing a STARlet automated liquid handler (Hamilton Company, Reno, NV). Paired-end sequencing (2×151bp) was performed using NovaSeq 6000 (Illumina, San Diego, CA) with a read depth of 94x and 123x and a genome length of 12288577 and 12267689 for isolates LNV001 (SRR23958537) and LNV002 (SRR23109153), respectively.

### Bioinformatic Analysis

The open-source software TheiaEuk was used to perform the *de novo* assembly, quality assessment, and genomic characterization of fungal genomes (21). Using the generated FASTa files, species taxon identification and clade typing was performed by Genomic Approximation Method for Bacterial Identification and Tracking (GAMBIT) (22). kSNP3 identified core genome single nucleotide polymorphisms (SNPs) between the two isolates with confirmation and gene identification performed by Snippy to the *C. auris* clade III reference strain B11221 (GCA_0022775015.1) (23, 24).

### Phenotypic Screens

#### Growth Curve

*C. auris* isolates were grown on CHROMagar plates at 36 °C for 48 hours and then a single colony was isolated and regrown for purity for another 48 hours at 36 °C. The high-throughput phenotypic screen was performed utilizing Biolog PM1 and PM2a phenotypic microarray plates (Biolog, Inc., Hayward, CA) per manufacturer’s protocol for *Saccharomyces cerevisiae*. Absorbance was taken every 6-8 hours for 72 hours. Graphs were designed in R studio with package growthcurver (25). Significance was determined by a student’s T-test of growth rate with Bonferroni method to adjust for a large data set.

#### Dilution Spot

Single *C. auris* colonies from respective isolates were grown on CHROMagar plates for mass growth at 36 °C for 48 hours. Both isolates were standardized by absorbance in RPMI broth. Ten-fold dilution series was performed to gain a final dilution of 10^−4^. Dilution spots were plated on RPMI agar and checked on every 24 hours for 72 hours.

### Sterol Analysis

Overnight cultures of *C. auris* strains were used to inoculate 20 mL MOPS buffered RPMI to 0.1 OD_590_. Cultures were grown for 18 hours at 35 °C, 170 rpm. Cells were then harvested and split into two samples, to enable determination of dry weight and extraction of sterols. An internal standard of 10 µg of cholestanol was added to each sample and sterols were extracted as previously described (26). Briefly, lipids were saponified using alcoholic KOH and non-saponifiable lipids extracted with hexane. Dried samples were derivatized by the addition of 0.1 mL BSTFA TMCS (99:1, Sigma) and 0.3 mL anhydrous pyridine (Sigma) and heating at 80 °C for 2 hours. TMS-derivatized sterols were analyzed and identified using gas chromatography-mass spectrometry (GC/MS) (Thermo 1300 GC coupled to a Thermo ISQ mass spectrometer, Thermo Scientific) and Xcalibur software (Thermo Scientific). The retention times and fragmentation spectra for known standards were used to identify sterols. Sterol composition was calculated from peak areas, as a mean of 3 replicates and the relative quantity of sterols present was determined using a standard curve of the internal standard (cholestanol) and lanosterol and the dry weight of the samples.

### EPIC Mediated ERG3 and ERG4 Modification in C. auris Isolate LNV002

#### EPIC Components

Episomal Plasmid Induced Cas9 (EPIC) mediated transformation of *C. auris* was performed as previously described (27, 28). Briefly, guide sequence primers targeting *ERG3* and *ERG4* were ligated into pJMR19 following Lgu1 (Thermo Scientific™) digestion as previously described (28). Transformation repair templates were amplified from gBlock sequences (Integrated DNA Technologies) by PCR using Phusion Green master mix manufacturer’s instructions (Thermo Scientific, Waltham, MA, USA) followed by subsequent QIAquick PCR purification (Qiagen). All strains, primers, and templates are listed in **Supplemental Table 1**.

#### *C. auris* Transformation

*C. auris* was cultured overnight to an OD_600_ of 1.8-2.2 and transformation reactions were assembled with salmon sperm DNA (Invitrogen), pJMR19, repair template DNA, and TE-LiAC + 55% PEG. EPIC positive transformants were selected by growth on nourseothricin (200 mg/L) supplemented YPD agar plates (28). Single colonies were patched onto YPD for screening. DNA isolation was achieved via treatment with a DNA extraction buffer consisting of 10 mM Tris pH 8.0, 2 mM EDTA, 0.2% Triton X-100, 200 ug/mL Proteinase K. *ERG3* and *ERG4* PCR amplification and subsequent Sanger sequencing (Hartwell Center, St. Jude Children’s Research Hospital) was used to screen positive transformants. Sequenced transformants were then replica plated for plasmid ejection as previously described and proper *ERG3, ERG4* sequence retention was confirmed with a second Sanger sequencing run (15, 27).

### Antifungal Susceptibility Testing

Minimum inhibitory concentrations (MIC) for amphotericin B (Sigma-Aldrich) were determined by broth microdilution in accordance with the M27-A4 from the Clinical Laboratory Standards Institute (CLSI) with considerations recommended by the CDC (14, 29). Amphotericin B MIC by E-test (BioMérieux, Marcy-l’Étoile, France) was determined per manufacturer’s instructions. All susceptibility testing was performed in biological triplicate and determined visually for growth inhibition at 24 hours.

## Results

### Patient Case Description

Consecutive *C. auris* isolates were collected over a two-month span in the summer of 2022 from a 76-year old female. Two isolates over the course of these two months were collected from the patient from the bronchial lavage and urine, respectively (**Table 1**). The initial isolate presented with a minimum inhibitory concentration (MIC) value of 0.19 µg/mL yielding susceptibility to amphotericin B while the latter isolate presented with a > 32 µg/mL MIC value yielding high resistance to amphotericin B (**Figure 1**,**2**).

**Table 1:** Case Study of Acquired Amphotericin B Resistance in *C. auris*.

| Isolate ID | Collection Date | Isolation Site | Amphotericin B MIC<br>($\mu\text{g/mL}$) | Number of Coding<br>Mutations |
| --- | --- | --- | --- | --- |
| LVN001 | July 2022 | Bronchial Lavage | 0.19 | 9 |
| LVN002 | September 2022 | Urine | >32 |  |

*C. auris* isolates were collected over a two-month span. Whole genome sequencing was performed on both isolates, revealing nine sequence differences that result in coding changes between the genomes.

**Figure 1:**
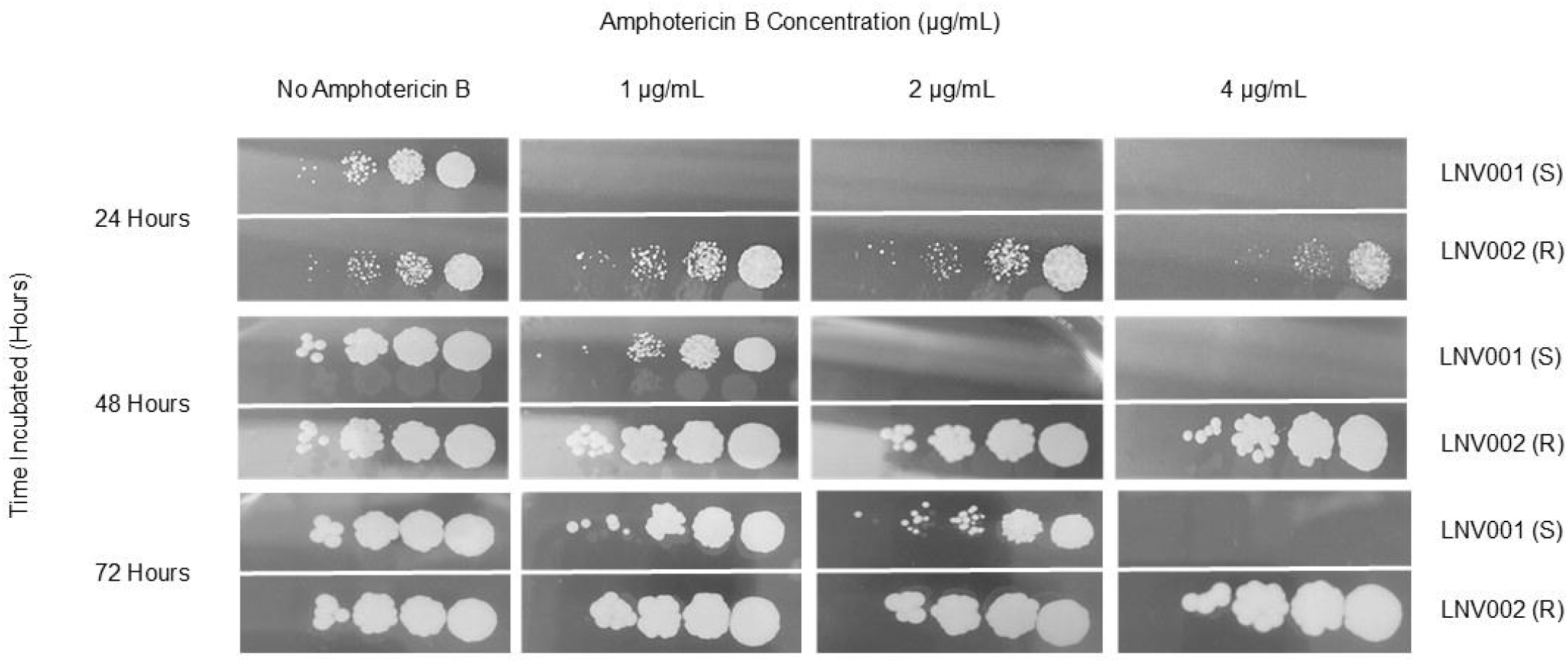
Dilution Spot Assay of Amphotericin B Susceptible and Resistant *C. auris* Isolates Ten-fold dilution series of LNV001 and LNV002 were plated on RPMI agar supplemented with varying concentration of amphotericin B and reading observed every 24 hours.

### Whole Genome Sequencing

The two isolates underwent paired-end sequencing on the NovaSeq 6000. Sequences were *de novo* assembled and quality checked by the bioinformatic workflow TheiaEuk (21).

Confirmation of the respective isolates as *C. auris* was performed using GAMBIT which further described each isolate as a member of Clade III (22). The sequences of the two isolates were genetically discordant from each other by nine sequence differences determined by the bioinformatic workflow Snippy. Nine mutations in coding regions were identified (**Table 2**) (24). Both isolates were resistant to fluconazole with a MIC value of > 256 µg/mL and each was found to possess a mutation encoding the VF125AL amino acid substitution in sterol 14α-demethylase encoded by *ERG11* which has been previously established to confer clinical fluconazole resistance and has exclusively been observed in Clade III isolates (7, 8). Between the amphotericin B susceptible and resistant isolates, the mutations of greatest interest to public health are the ten base pair deletion causing a frameshift at amino acid 71 leading to a premature stop codon at amino acid 261 in the C-5 sterol desaturase gene, *ERG3*, and the nonsense mutation causing a premature stop codon at amino acid 106 in the delta (24 (24(1)))-sterol reductase gene, *ERG4*. Since amphotericin B is hypothesized to exert antifungal activity through direct interaction with ergosterol we focused on *ERG3* and *ERG4* of the ergosterol biosynthesis pathway.

**Table 2:** SNPs Differences Between LNV001 and LNV002 in Coding Regions.

| Protein (Gene) | Predicted Ortholog | Gene Locus | Mutation | Mutation Type |
| --- | --- | --- | --- | --- |
| **C-5 sterol desaturase**<br>**(*ERG3*)** | **C-5 sterol desaturase (*ERG3*)** | **CJI97_003811** | **G71fs -> S261\*** | **Frameshift/Stop** |
| **Delta (24 (24(1)))-sterol**<br>**reductase (*ERG4*)** | **Delta (24 (24(1)))-sterol**<br>**reductase (*ERG4*)** | **CJI97_002908** | **L106\*** | **Stop** |
| 1-phosphatidylinositol 4-kinase | 1-phosphatidylinositol 4-kinase | CJI97_001150 | N126D | Missense |
| Retromer subunit VPS35 | Retromer subunit VPS35 | CJI97_003028 | I175V | Missense |
| Uncharacterized protein | Major Facilitator Superfamily | CJI97_000798 | F232S | Missense |
| Uncharacterized protein | Mediator of RNA polymerase II<br>transcription subunit 5 | CJI97_002146 | F275fs | Frameshift |
| Uncharacterized protein | Selenoprotein O | CJI97_002594 | R243fs | Frameshift |
| Uncharacterized protein | *STE3* | CJI97_005168 | A178V | Missense |
| Uncharacterized protein | Transcriptional regulator *ADR1* | CJI97_002218 | E41Q | Missense |

Nine coding regions have been identified to possess mutations from the former susceptible isolate. *ERG3* and *ERG4* which are hypothesized to be responsible for the resistant phenotype are bolded. Gene loci are from the clade III reference strain B11221.

### EPIC Mediated Reversion of ERG3 and ERG4 to Wildtype

To identify the influence of each mutated gene on the amphotericin B resistance, we utilized the EPIC genetic manipulation system to revert the *ERG3* and *ERG4* sequences to wildtype (matching LNV001 and the B11221 reference) in LNV002 (28). While we were able to generate two independent *ERG3*^*WT*^ single reversions in LNV002, we were unable to generate *ERG4*^*WT*^ single reversions in LNV002. However, once *ERG3* had been reverted to wildtype, we were able to generate two independent *ERG3*^WT^, *ERG4*^WT^ double correction strains. When testing amphotericin B susceptibility using the diffusion test strip (E-test, bioMérieux) method, as recommended by the CDC, the *ERG3*^WT^ reversion in LNV002 resulted in dramatically increased amphotericin B susceptibility with a MIC shift from >32 mg/L to 0.38 mg/L (**Figure 2A**).

**Figure 2:**
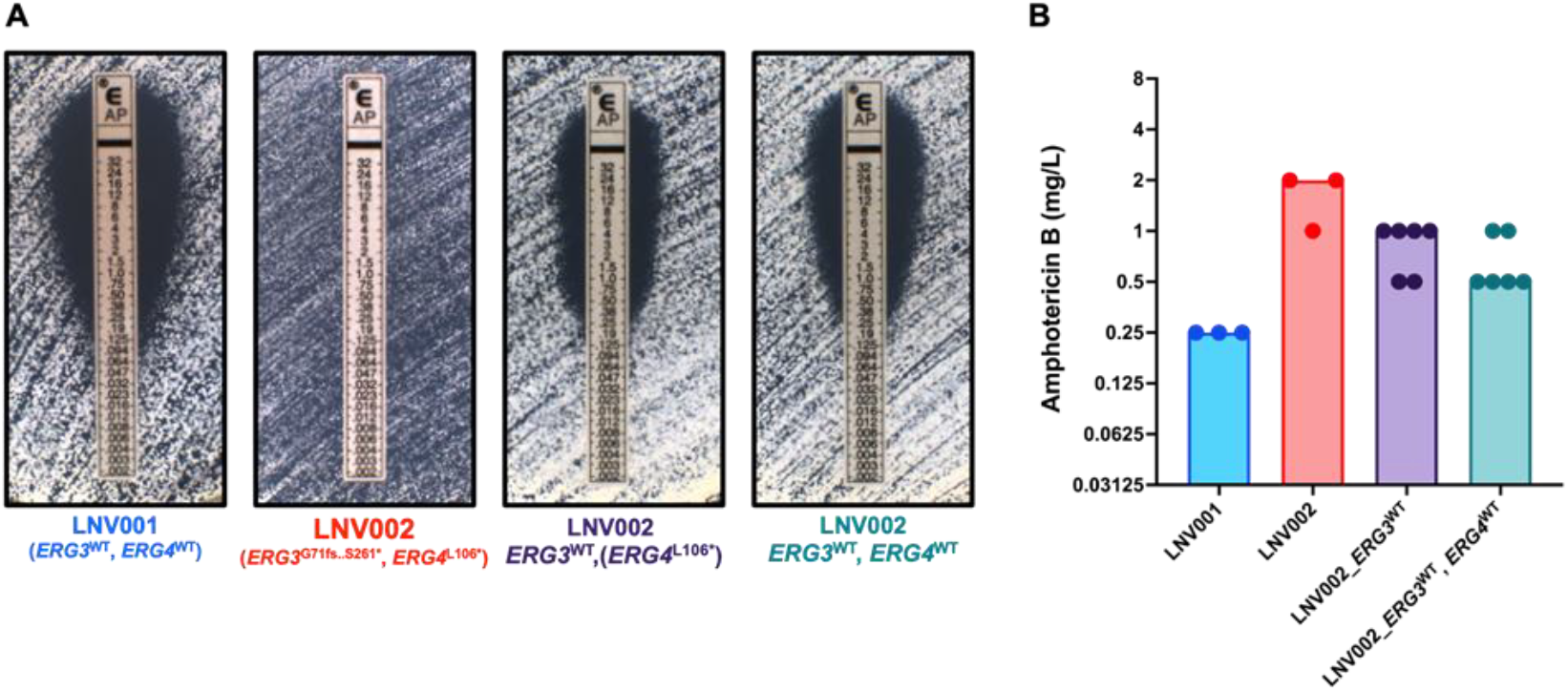
Amphotericin B Minimum Inhibitory Concentrations following *ERG3* and *ERG4* reversion to wildtype. A) Representative images of Amphotericin B Minimum Inhibitory Concentrations (MICs) at 24 hours as determined by E-tests (bioMérieux). B) MICs determined by broth microdilution in accordance with CLSI susceptibility testing. MICs were read visually for 100% growth inhibition at 24 hours. Bars represent the modal MIC with points plotted for three biological replicates for each isolate and independent strain with MIC values for two independently derived LNV002_ERG3^WT^ and LNV002_ERG3^WT^, ERG4^WT^ strains shown.

However, the additional reversion of *ERG4* to the wildtype allele (in strain LNV002_*ERG3*^WT^, *ERG4*^WT^) did not further increase amphotericin B susceptibility. By comparison, when testing amphotericin B MIC using the broth microdilution method, reversion of the *ERG3* gene to the wildtype sequence resulted in a more modest decrease in amphotericin B MIC, and correction of both *ERG3* and *ERG4* to the wildtype allele resulted in a further one-dilution decrease in amphotericin B MIC (**Figure 2B**).

### Absence of Ergosterol

We sought to determine if LNV002 produced ergosterol via sterol profiling because of the predicted interaction between amphotericin B and ergosterol. TMS-derivatized sterols in biological triplicate were analyzed and identified using GC/MS and Xcalibur software. The sterol profile of the susceptible LNV001 isolate largely consists of ergosterol and lanosterol (**Table 3, Supplemental Figure 1**). The sterol profile of the resistant isolate (LNV002) is mainly composed of ergosta-7,22,24(28)-trienol, episterol, and lanosterol. Ergosterol was not detectable in the resistant isolate.

**Table 3:** Sterol Composition of LNV001 and LNV002.

| Sterol | Percentage of Total Sterol |  |  |  |
| --- | --- | --- | --- | --- |
|  | LNV001 (S) |  | LNV002 (R) |  |
|  | Mean | ±Standard Deviation | Mean | ±Standard Deviation |
| Ergosta-5,8,22,24(28)-tetraenol | 0.3 | 0.1 |  |  |
| Ergosta-5,8,22-trienol | 0.1 | 0.0 |  |  |
| Ergosta-8,22,24(28)-trienol |  |  | 1.5 | 0.0 |
| Zymosterol | 0.3 | 0.0 |  |  |
| **Ergosterol** | **45.2** | **6.4** |  |  |
| Ergosta 7,22 dienol | 0.6 | 0.1 |  |  |
| unidentified sterol |  |  | 1.1 | 0.1 |
| 4,14-dimethyl zymosterol | 0.3 | 0.1 | 2.8 | 1.0 |
| Fecosterol (Ergosta-8,24(28)-dienol) |  |  | 0.4 | 0.3 |
| **Ergosta-7,22,24(28)-trienol** |  |  | **53.7** | **8.6** |
| 14-methyl fecosterol | 2.0 | 0.5 | 3.2 | 1.0 |
| Ergosta-5,7-dienol | 5.7 | 7.6 |  |  |
| unidentified sterol |  |  | 0.6 | 0.1 |
| **Episterol (Ergosta-7,24(28)-dienol)** |  |  | **17.3** | **4.8** |
| Ergosta-7-enol | 0.3 | 0.2 |  |  |
| 4,4 dimethyl-ergosta 8,14,24(28)-trienol | 0.2 | 0.0 | 0.1 | 0.0 |
| 14-methyl ergosta-8,24(28)-dien-3-6-diol | 0.2 | 0.0 |  |  |
| **Lanosterol** | **39.7** | **1.3** | **17.3** | **2.1** |
| 4-methyl ergosta-8,24(28)-dienol | 0.7 | 0.5 |  |  |
| 4,4-dimethyl cholesta-8,24-dienol | 1.4 | 0.7 | 0.8 | 0.2 |
| 4,4-dimethyl cholesta-8,14,24-dienol |  |  | 0.3 | 0.1 |
| Eburicol | 3.0 | 0.5 | 0.9 | 0.3 |
| **Total** | **100.0** |  | **100.0** |  |

TMS-derivatized sterols were analyzed and identified using GC/MS and XCALIBUR software.
Sterols that mainly make up each isolate are bolded. Sterol compositions are out of 100%.

### Fitness Cost Associated with Amphotericin B Non-Susceptibility

Amphotericin B resistance has been observed rarely for most *Candida* species (30). Evidence suggests that this is due to severe fitness defects caused by mutations in the ergosterol biosynthesis pathway (31). Since LNV002 had mutations in the ergosterol biosynthesis pathway, we hypothesized that there could be a fitness cost associated with *ERG3* and *ERG4* mutations in *C. auris*. To test this, we performed growth assays over 72 hours on 190 different carbon sources. We determined fitness differences between LNV001 and LNV002 as described in the materials and methods (**Supplemental Table 2**). Fourteen carbon sources were determined to be statistically significant for fitness defects in isolate LNV002. Carbon sources β-D-Allose, Phenylethylamine, Xylitol, α-Keto-Glutaric Acid, Glycyl-L Glutamic Acid, and Glycl-L-Aspartic Acid are associated with the highest fitness defects (**Figure 3**). In general, this experiment concluded that there is a fitness cost associated with the nine coding mutations present in the resistant isolate when utilizing certain carbon sources for energy.

**Figure 3:**
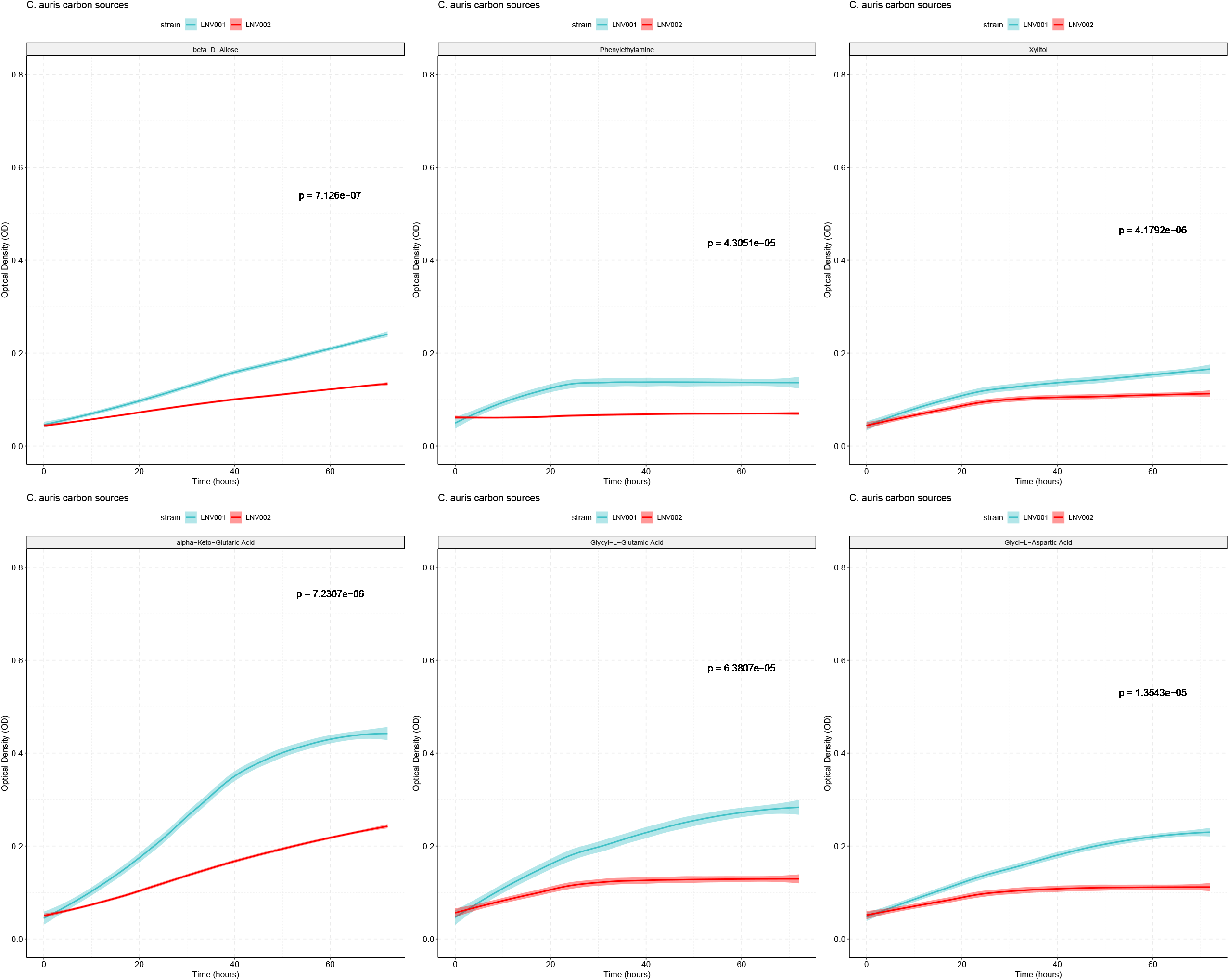
Significant Growth of Amphotericin B Susceptible Isolate Compared to Resistant Isolate Biolog Phenotypic plates PM1 and PM2a were inoculated with LNV001 and LNV002. Optical density was measured every 6-8 hours for 72 hours. Blue line is LNV001 and red line is LNV002. Standard deviation is denoted by the blurring surrounding the lines. Significance determined by a student’s T-test with the Bonferroni correction applied (α = 0.000263).

## Discussion

### Amphotericin B is Critical to Managing this Public Health Threat

Over 90% of *C. auris* clinical isolates demonstrate resistance to fluconazole increasing our dependence on echinocandins and amphotericin B as the recommended frontline therapy for the treatment of invasive candidiasis and infections caused by *C. auris* specifically (14, 32). In 2021, the CDC noted that the number of echinocandin-resistant *C. auris* infections have tripled since 2020, leaving amphotericin B as a last resort to potentially treat these cases (3). Even though it has been observed that approximately 30% of *C. auris* isolates are resistant to amphotericin B, this agent remains a crucial drug in the antifungal armamentarium due to its broad-spectrum antifungal activity (10). *C. auris* is an emerging public health threat, and understanding its genetic basis of resistance is crucial for effective infection prevention and control.

### Interpretation of CRISPR Results

From a clinical case, we isolated a pair of patient isolates, one susceptible and one resistant to amphotericin B collected within two months of each other. Between both isolates, a narrow list of non-synonymous mutations was revealed. Due to the involvement of ergosterol in the mechanism of action of amphotericin B, we focused our efforts on mutations present in genes which are predicted to play a role in the ergosterol biosynthesis pathway, *ERG3* and *ERG4*.

Correction of the *ERG3* frameshift mutation (*ERG3*^G71fs..S261*^) in LNV002 restored clinical amphotericin B susceptibility, thus demonstrating a documented case of acquired amphotericin B resistance resulting from a deleterious *ERG3* mutation in a clinical *C. auris* isolate. The restored LNV002_*ERG3*^*WT*^ strain was found to possess a MIC value below the current determination of clinical resistance set by the CDC of 2 µg/mL. When *ERG3* and *ERG4* were both restored to the wildtype sequence in LNV002, the MICs remained relatively unchanged from that of the individual LNV002_*ERG3*^WT^ strain, leaving only a small susceptibility difference persisting between LNV002_*ERG3*^WT^, *ERG4*^WT^ and that of LNV001. It remains possible that one of the other identified mutations observed in LNV002 may modestly contribute to amphotericin B resistance and account for the 0.5 to 1-dilution difference in MIC not attributable to mutations in *ERG3* or *ERG4*. One mutation of note that we speculate as a potential gene modifier candidate is gene locus CJI97_002218, an ortholog to *ADR1*, a transcriptional regulator in *C. albicans*. It has been found that *ADR1* has undergone transcriptional rewiring from *S. cerevisiae* where now it directs the regulation of the ergosterol biosynthesis pathway in *C. albicans* (33). It is plausible that *ADR1* in *C. auris* could also contribute to the regulation of the ergosterol biosynthesis pathway, making this missense mutation a primary candidate for the difference in MIC between the LNV002_*ERG3*^WT^, *ERG4*^WT^ corrected strains and the susceptible isolate LNV001.

Additionally, the small susceptibility difference could be attributed to methodological testing differences when determining MIC. Microbroth dilution determines MIC by growth inhibition in broth while E-test diffusion strips determine MIC by an ellipse on solid media. Specifically for amphotericin B, the dynamic testing range for microbroth dilutions has been observed to be highly condensed which leaves more variability in interpretation of growth differences. This observation of testing range combined with differences between broth and solid media are the likely factors contributing to the observed MIC difference. Nevertheless, both microbroth dilution and E-test diffusion strips demonstrate a consistent restoration of amphotericin B susceptibility upon *ERG3* mutation correction.

It is also notable that within the boundaries of our methods, the generation of a LNV002_*ERG4*^*WT*^ independent strain was not found to be possible. Therefore, we hypothesize that the *ERG3* mutation may have occurred prior to the *ERG4* mutation with the subsequent deleterious *ERG4* mutation potentially compensating fitness loss associated with amphotericin B resistance. Further elucidation into the role of *ERG4* in this process will require additional study.

### Replacement of Ergosterol

In fungal cell membranes, ergosterol is a vital component responsible for membrane fluidity regulation, making the ergosterol biosynthesis pathway an attractive target for conventional antifungal use and drug development. In *C. albicans* and *S. cerevisiae* the loss of function of lanosterol 14α-demethylase and C-5 sterol desaturase (*ERG11* and *ERG3*) have been associated with acquired amphotericin B resistance and for the exchange of ergosterol for other sterols such as lanosterol, eburicol, and 4,14-dimethyl-zymosterol in the cell membrane (34, 35). The resistant isolate described here does not produce ergosterol and showed enrichment of lanosterol, episterol, and late-stage sterols such as ergosta-7,22,24(28)-trienol. This altered membrane composition which likely leads to amphotericin B resistance is consistent with a loss of function of *ERG3* and *ERG4* based on their function in ergosterol biosynthesis. We hypothesize that these cells are able to survive because late-stage sterols are proposed to be able to stabilize the cell membrane in the absence of ergosterol.

### Fitness Cost Identified with Amphotericin B Resistant Isolate

Amphotericin B resistance is rarely observed for most *Candida spp*. This may be due to the fitness defects of mutations in the ergosterol biosynthesis pathway (31). In characterizing this resistant phenotype, a comparison between the susceptible and resistant isolates supplemented with 190 different carbon sources revealed a statistically significant fitness defect present on a few of these sources. This data may indicate a narrow fitness cost for combinatorial mutations in *ERG3* and *ERG4* in this isolate. Conversely, it is possible that among the other seven mutations detected by whole genome sequencing there are one or more modifiers that alleviate the fitness cost associated with perturbation of the ergosterol biosynthesis pathway.

## Conclusion

*C. auris* is spreading broadly and the initially identified four major clades are no longer endemic solely to the area in which they were initially associated (36). The spread of this agent is resulting in extensive morbidity and mortality to patients in health care facilities across the world. Antifungal drug resistance will continue to evolve and spread unless we can identify and implement effective infection control mechanisms. With only three primary anti-fungal drug classes available to treat fungal infections in the U.S. and the multi-drug resistant characteristics of *C. auris*, the emphasis on anti-microbial stewardship and along with the development of novel antifungals are more essential than ever to combat this pathogen.

This article characterizes a clinical case description of *C. auris* amphotericin B resistance resulting from a frameshift mutation in *ERG3*. In some outbreaks, one third of *C. auris* isolates demonstrate amphotericin B resistance that is often attributed to an unidentified mechanism (9). This finding represents a significant advancement in understanding antifungal resistance in *C. auris* and the knowledge generated can be used to design diagnostic tests to combat antifungal drug failure.

## Acknowledgments

This research was supported in part by ALSAC and the National Cancer Institute grant P30 CA021765, by NIH NIAID grant R01 AI169066 awarded to P.D.R. and C.A.C, by NIAID grant U19AI110818 to the Broad Institute (C.A.C.), by the St. Jude Children’s Research Hospital Children’s Infection Defense Center grant (J.M.R.) and the Society of Infectious Diseases Pharmacists Young Investigator Research Award granted to J.M.R.. This publication was supported by the Nevada State Department of Health and Human Services through Grant # 5 NU50CK000560-05-00 from Epidemiology and Laboratory Capacity for Infectious Diseases (ELC). Its contents are solely the responsibility of the authors and do not necessarily represent the official views of the Department nor Epidemiology and Laboratory Capacity for Infectious Diseases (ELC).

## Notes

### Competing Interest Statement

The authors have declared no competing interest.

