## Supplemental Figure 1 and Supplemental Tables 1 & 2 for "Acquired Amphotericin B Resistance Attributed to a Mutated *ERG3* in *Candidozyma auris*"

### Slide 1
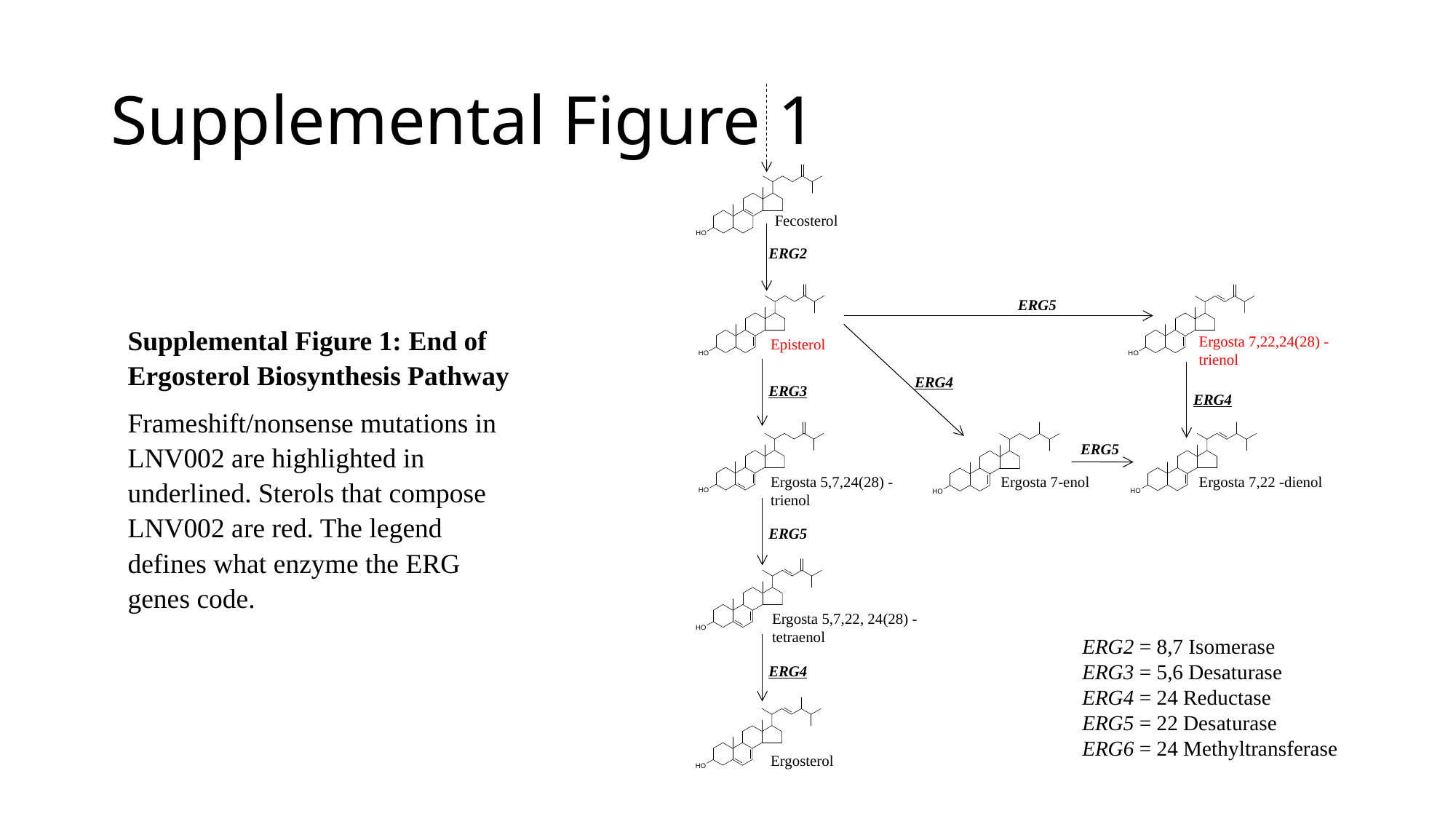

#### Supplemental Figure 1
Fecosterol
ERG2
ERG5
Ergosta 7,22,24(28) -trienol
Episterol
ERG4
ERG3
ERG4
Ergosta 5,7,24(28) -trienol
Ergosta 7-enol
Ergosta 7,22 -dienol
ERG5
Ergosta 5,7,22, 24(28) -tetraenol
ERG4
Ergosterol
ERG5
Supplemental Figure 1: End of Ergosterol Biosynthesis Pathway
Frameshift/nonsense mutations in LNV002 are highlighted in underlined. Sterols that compose LNV002 are red. The legend defines what enzyme the ERG genes code.
ERG2 = 8,7 Isomerase
ERG3 = 5,6 Desaturase
ERG4 = 24 Reductase
ERG5 = 22 Desaturase
ERG6 = 24 Methyltransferase

### Slide 2
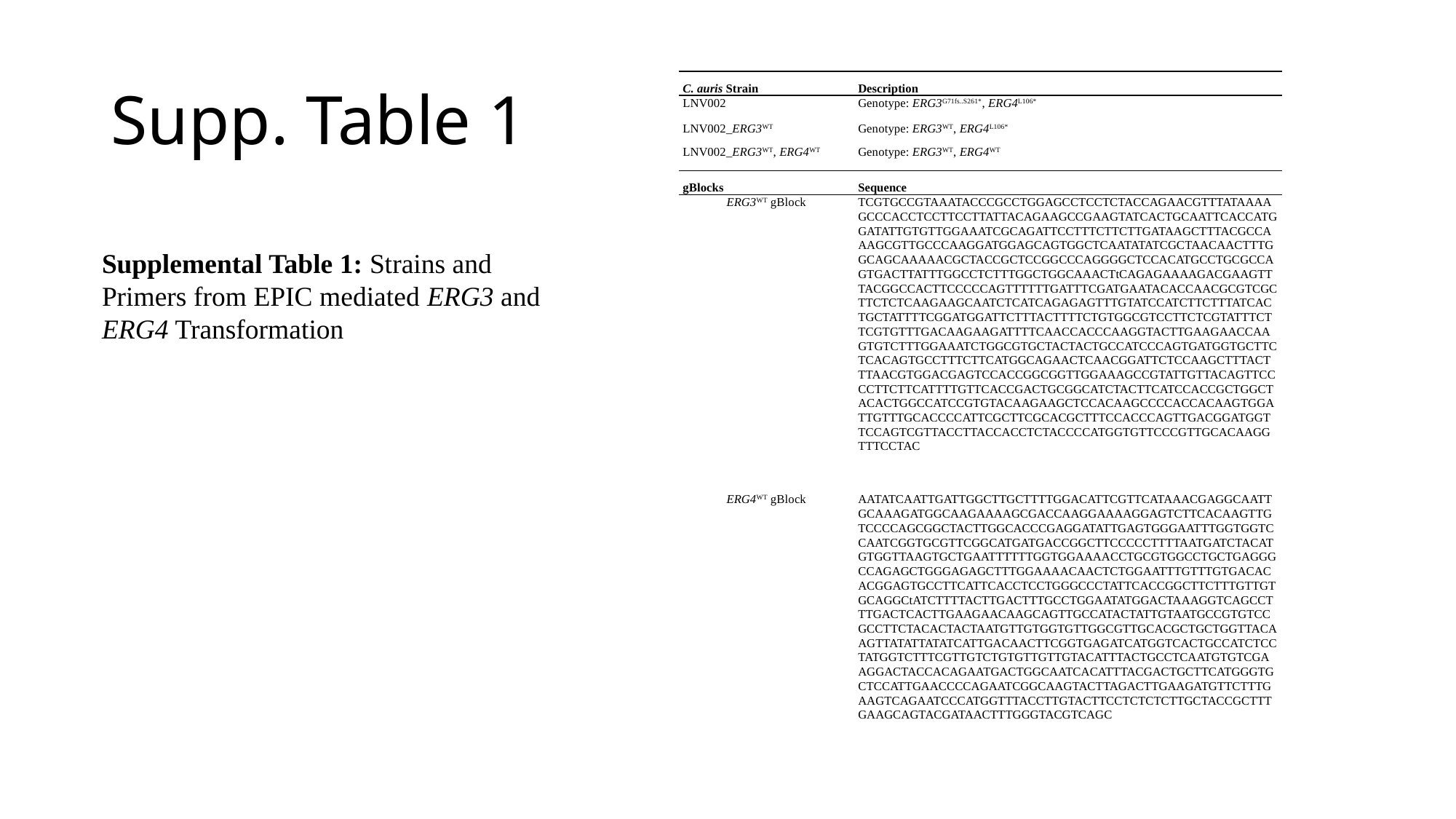

#### Supp. Table 1
| C. auris Strain | Description |
| --- | --- |
| LNV002 | Genotype: ERG3^G71fs..S261\*^, ERG4^L106\*^ |
| LNV002\_ERG3^WT^ | Genotype: ERG3^WT^, ERG4^L106\*^ |
| LNV002\_ERG3^WT^, ERG4^WT^ | Genotype: ERG3^WT^, ERG4^WT^ |
| gBlocks | Sequence |
| ERG3^WT^ gBlock | TCGTGCCGTAAATACCCGCCTGGAGCCTCCTCTACCAGAACGTTTATAAAAGCCCACCTCCTTCCTTATTACAGAAGCCGAAGTATCACTGCAATTCACCATGGATATTGTGTTGGAAATCGCAGATTCCTTTCTTCTTGATAAGCTTTACGCCAAAGCGTTGCCCAAGGATGGAGCAGTGGCTCAATATATCGCTAACAACTTTGGCAGCAAAAACGCTACCGCTCCGGCCCAGGGGCTCCACATGCCTGCGCCAGTGACTTATTTGGCCTCTTTGGCTGGCAAACTtCAGAGAAAAGACGAAGTTTACGGCCACTTCCCCCAGTTTTTTGATTTCGATGAATACACCAACGCGTCGCTTCTCTCAAGAAGCAATCTCATCAGAGAGTTTGTATCCATCTTCTTTATCACTGCTATTTTCGGATGGATTCTTTACTTTTCTGTGGCGTCCTTCTCGTATTTCTTCGTGTTTGACAAGAAGATTTTCAACCACCCAAGGTACTTGAAGAACCAAGTGTCTTTGGAAATCTGGCGTGCTACTACTGCCATCCCAGTGATGGTGCTTCTCACAGTGCCTTTCTTCATGGCAGAACTCAACGGATTCTCCAAGCTTTACTTTAACGTGGACGAGTCCACCGGCGGTTGGAAAGCCGTATTGTTACAGTTCCCCTTCTTCATTTTGTTCACCGACTGCGGCATCTACTTCATCCACCGCTGGCTACACTGGCCATCCGTGTACAAGAAGCTCCACAAGCCCCACCACAAGTGGATTGTTTGCACCCCATTCGCTTCGCACGCTTTCCACCCAGTTGACGGATGGTTCCAGTCGTTACCTTACCACCTCTACCCCATGGTGTTCCCGTTGCACAAGGTTTCCTAC |
| ERG4^WT^ gBlock | AATATCAATTGATTGGCTTGCTTTTGGACATTCGTTCATAAACGAGGCAATTGCAAAGATGGCAAGAAAAGCGACCAAGGAAAAGGAGTCTTCACAAGTTGTCCCCAGCGGCTACTTGGCACCCGAGGATATTGAGTGGGAATTTGGTGGTCCAATCGGTGCGTTCGGCATGATGACCGGCTTCCCCCTTTTAATGATCTACATGTGGTTAAGTGCTGAATTTTTTGGTGGAAAACCTGCGTGGCCTGCTGAGGGCCAGAGCTGGGAGAGCTTTGGAAAACAACTCTGGAATTTGTTTGTGACACACGGAGTGCCTTCATTCACCTCCTGGGCCCTATTCACCGGCTTCTTTGTTGTGCAGGCtATCTTTTACTTGACTTTGCCTGGAATATGGACTAAAGGTCAGCCTTTGACTCACTTGAAGAACAAGCAGTTGCCATACTATTGTAATGCCGTGTCCGCCTTCTACACTACTAATGTTGTGGTGTTGGCGTTGCACGCTGCTGGTTACAAGTTATATTATATCATTGACAACTTCGGTGAGATCATGGTCACTGCCATCTCCTATGGTCTTTCGTTGTCTGTGTTGTTGTACATTTACTGCCTCAATGTGTCGAAGGACTACCACAGAATGACTGGCAATCACATTTACGACTGCTTCATGGGTGCTCCATTGAACCCCAGAATCGGCAAGTACTTAGACTTGAAGATGTTCTTTGAAGTCAGAATCCCATGGTTTACCTTGTACTTCCTCTCTCTTGCTACCGCTTTGAAGCAGTACGATAACTTTGGGTACGTCAGC |
Supplemental Table 1: Strains and Primers from EPIC mediated ERG3 and ERG4 Transformation

### Slide 3
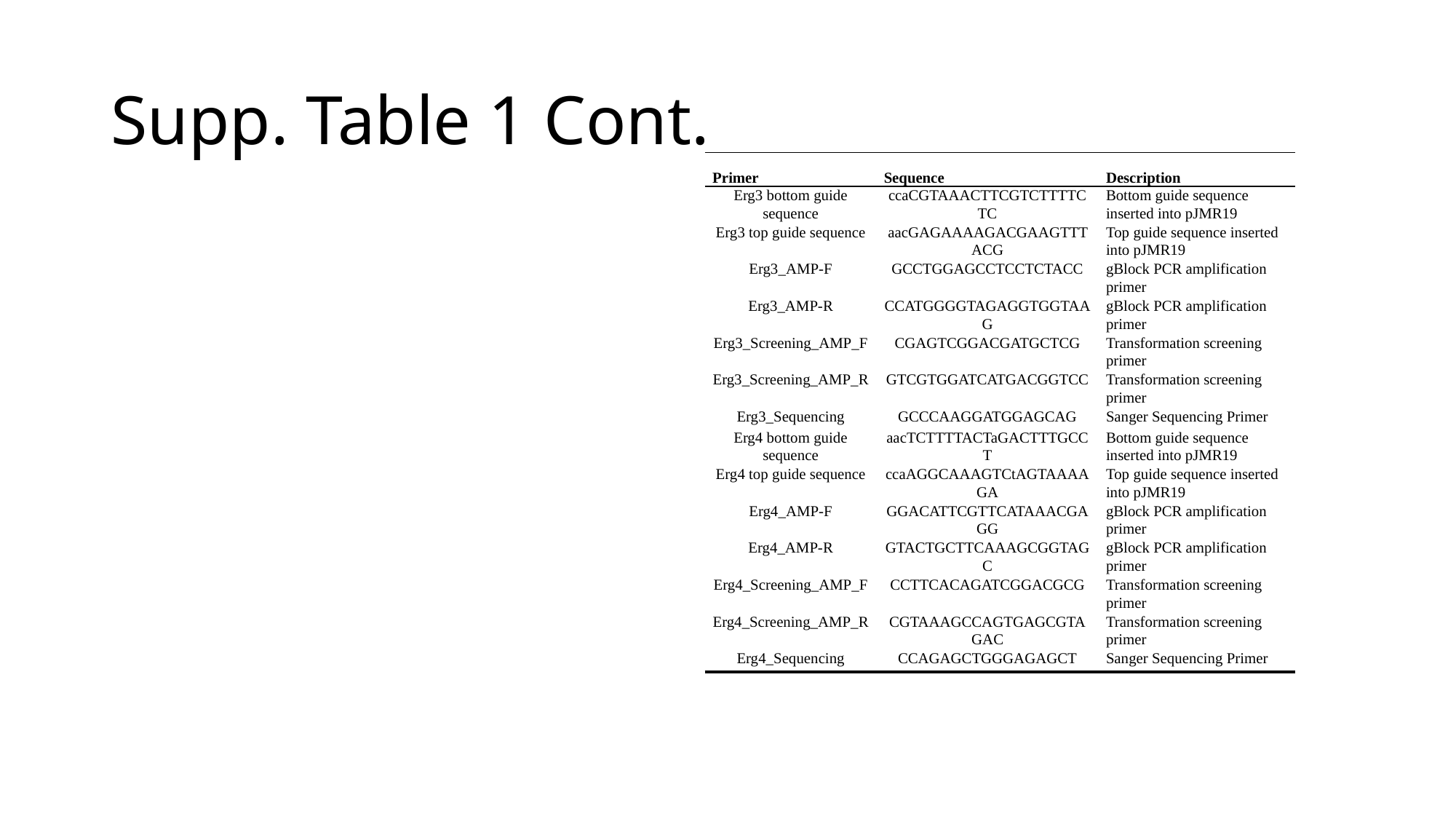

#### Supp. Table 1 Cont.
| Primer | Sequence | Description |
| --- | --- | --- |
| Erg3 bottom guide sequence | ccaCGTAAACTTCGTCTTTTCTC | Bottom guide sequence inserted into pJMR19 |
| Erg3 top guide sequence | aacGAGAAAAGACGAAGTTTACG | Top guide sequence inserted into pJMR19 |
| Erg3\_AMP-F | GCCTGGAGCCTCCTCTACC | gBlock PCR amplification primer |
| Erg3\_AMP-R | CCATGGGGTAGAGGTGGTAAG | gBlock PCR amplification primer |
| Erg3\_Screening\_AMP\_F | CGAGTCGGACGATGCTCG | Transformation screening primer |
| Erg3\_Screening\_AMP\_R | GTCGTGGATCATGACGGTCC | Transformation screening primer |
| Erg3\_Sequencing | GCCCAAGGATGGAGCAG | Sanger Sequencing Primer |
| Erg4 bottom guide sequence | aacTCTTTTACTaGACTTTGCCT | Bottom guide sequence inserted into pJMR19 |
| Erg4 top guide sequence | ccaAGGCAAAGTCtAGTAAAAGA | Top guide sequence inserted into pJMR19 |
| Erg4\_AMP-F | GGACATTCGTTCATAAACGAGG | gBlock PCR amplification primer |
| Erg4\_AMP-R | GTACTGCTTCAAAGCGGTAGC | gBlock PCR amplification primer |
| Erg4\_Screening\_AMP\_F | CCTTCACAGATCGGACGCG | Transformation screening primer |
| Erg4\_Screening\_AMP\_R | CGTAAAGCCAGTGAGCGTAGAC | Transformation screening primer |
| Erg4\_Sequencing | CCAGAGCTGGGAGAGCT | Sanger Sequencing Primer |

### Slide 4
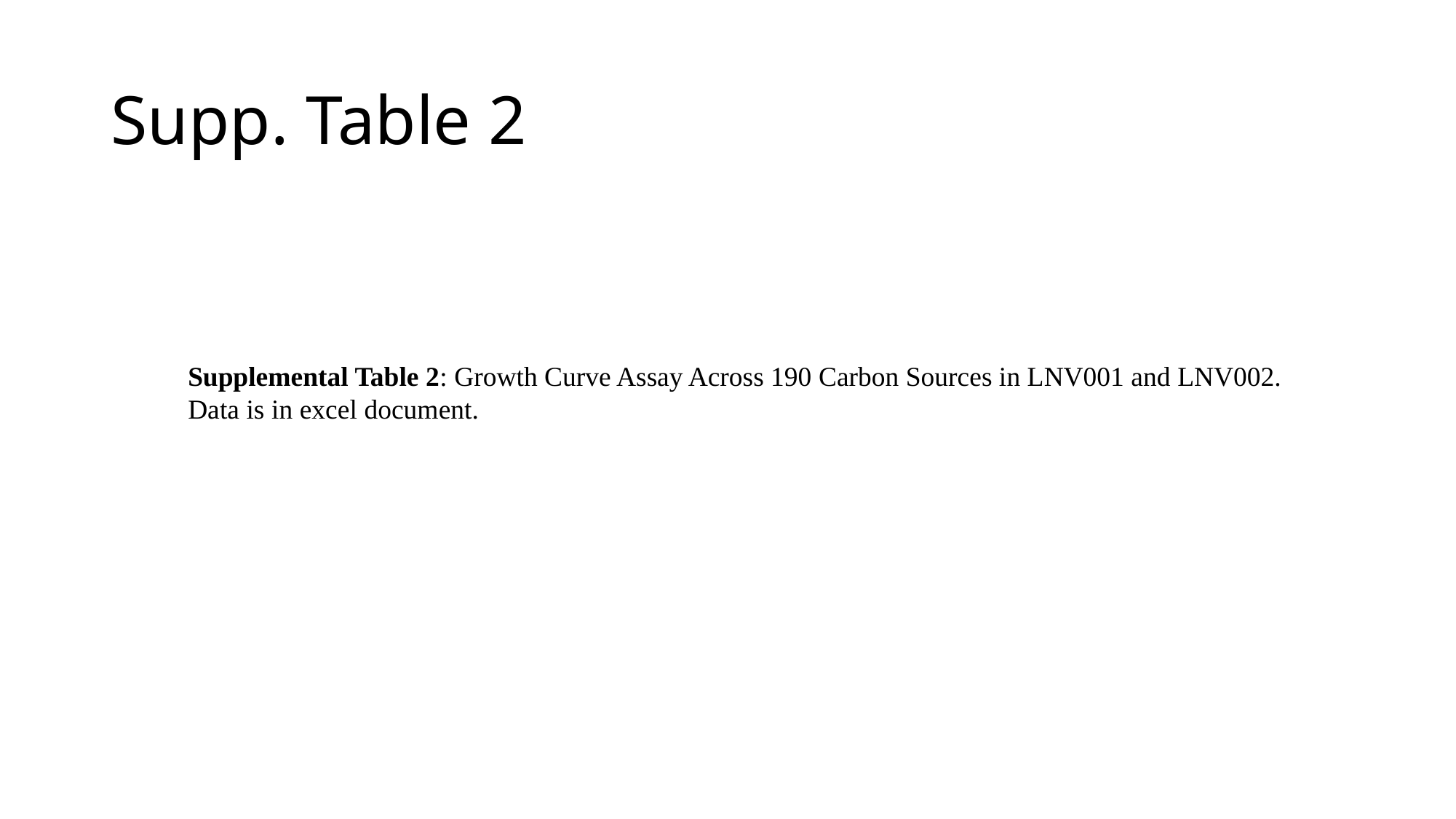

#### Supp. Table 2
Supplemental Table 2: Growth Curve Assay Across 190 Carbon Sources in LNV001 and LNV002.
Data is in excel document.
